# Transcription Profile And Pathway Analysis Of The Endocannabinoid Receptor Inverse Agonist AM630 In The Core And Infiltrative Boundary Of Human Glioblastoma Cells

**DOI:** 10.1101/2021.12.29.474430

**Authors:** Gareth Williams, David Chambers, Ruman Rahman, Francisco Molina-Holgado

## Abstract

**Background:** We have previously reported that the endocannabinoid receptor inverse agonist AM630 is a potent inhibitor of isocitrade dehydrogenase-1 wild-type glioblastoma (GBM) core tumor cell proliferation. To uncover the mechanism behind the anti-tumour effects we have performed a transcriptional analysis of AM630 activity both in the tumour core cells (U87) and the invasive margin cells (GIN-8), the latter representing a better proxy of post-surgical residual disease.

**Results:** The core and invasive margin cells exhibited markedly different gene expression profiles and only the core cells had high expression of a potential AM630 target, the CB1 receptor. Both cell types had moderate expression of the HTR2B serotonin receptor, a reported AM630 target. We found that the AM630 driven transcriptional response was substantially higher in the central cells than in the invasive margin cells, with the former driving the up regulation of immune response and the down regulation of cell cycle and metastatic pathways and correlating with transcriptional responses driven by established anti-neoplastics as well as serotonin receptor antagonists.

**Conclusion:** Our results highlight the different responsiveness of the core and invasive margin cells. Taken together, whilst our findings identify AM630 as an anti-neoplastic drug, showing a high correlation with known anti-proliferative drugs, we find distinct drug sensitivies of the infiltrative margin relative to contrast-enhanced core regions of GBM upon which failed molecular targeted therapies to date have been predicated.

## Introduction

Cell fate decisions are key in the homeostatic maintenance of the cellular *milieu* as well as in the survival of tissues and organisms. An uncontrolled cell proliferation occurs in tumoral cancer cells. This is a direct consequence of dysregulated cell cycle phases due to a lack of effective cell cycle DNA check points as the cell grows and divides. In the central nervous system (CNS), different systems control cell fate decisions to maintain an effective and functional brain circuitry[1]. Pathological alterations of this brain cellular network occur in different pathologies such as brain tumours[2].

The high-grade malignant brain tumour, isocitrate dehydrogenase-1 wild-type glioblastoma (GBM), is the most frequent and aggressive primary *de novo* tumour of the CNS, with a median survival of 14.6 months from diagnosis in patients multimodally treated with surgery, radiotherapy, and chemotherapy[3]. Despite multiple clinical trials and studies from several laboratories worldwide, there is no cure, unlike the treatment scenario for other tumours[4]. Median survival after diagnosis is virtually unchanged since records began in the 1930s, with extent of surgical resection being the best indicator of survival[5].

In the developed brain the neuromodulatory activity of the endocannabinoid (eCB) system is the control of neural excitability through retrograde signaling to inhibit presynaptic transmitter release[6]. However, during development, the eCB system functions to promote neurite outgrowth[7]. Pro-proliferative activity associated with eCB signaling has been reported in the neural stem cell niche of the subventricular zone and hippocampus of rodents[8]. RNAseq analysis of brain cell types reveal potential eCB responsiveness of microglia, oligodendrocytes, and astrocytes[9]. These effects are key in the neuroimmune and neuroprotective interactions of the eCB in response to different insults. In addition, the eCB network is an important regulator of brain cell fate determination (i.e. proliferation, migration and differentiation) in healthy and in pathological conditions. Signal transduction *via* cannabinoid receptors (CB1 and CB2) or *via* orphan GPCRs cannabinoid receptor-like receptors (GPR18, GPR55 and GPR119), leads to the proliferation, differentiation, and cell death events of brain cells, and this has important consequences for neural development and brain repair[10]. Thus, associated signalling pathways of the brain cannabinoid system are known to mediate several events in both the developing and adult nervous systems[11, 12]. The eCB system has also been widely studied in the context of cancer[11] with overexpression of eCBs and their receptors associated with tumour aggressiveness[13]. A dysregulation of eCB levels, which produced a modified responsiveness to specific ligands, has been shown in different cancer cell lines[14]. Interestingly, the peripheral CB receptor CB2 has been characterized as a novel murine proto-oncogene and confirmed to have a role in leukaemia development[15] and in the promotion of renal cell carcinoma prognosis and progression[16]. Other studies have implicated CB2 receptors as regulators of HER2 pro-oncogenic signalling, demonstrating that genetic inactivation of the CB2 receptor impairs tumour generation and progression in MMTV-neu mice[17]. However, the precise molecular mechanisms directing many of the above cellular events are still far from being completely understood.

In previous studies, we have found that the pharmacological blockade of the CB1 or CB2 receptor signaling pathways impairs (by targeting the mitochondrial unfold protein response (UPRmt)) the *in vitro* proliferation of human GBM cells obtained from the core region (U87) suggesting that CB2 cannabinoid receptors are somehow involved in the proliferation of these GBM cells[18]. The *in vivo* or *in vitro* effects of eCBs in tumour cell fate are an open debate in the scientific community that needs deeper investigation as the studies are limited and the molecular mechanisms underlying eCB activity in this context are poorly understood. Moreover, there is an increasing appreciation of GBM inter- and intra-tumour heterogeniety which manifests via evolutionary mechanisms[19], and we have shown that the infiltrative margin of GBM exhibits distinct transcriptomic profiles from other intra-tumour regions and is more representative of post-surgical minimal volume residual disease[20].

With the aim of uncovering the biological mechanisms behind the anti-tumour activity associated with eCB receptor inhibition we performed a detailed investigation into the gene expression changes driven by receptor inhibition in the context of the core and invasive margin cell populations of GBM. Specifically, we performed a microarray analyses of the gene expression perturbation driven by the CB2 receptor inverse agonist AM630 in GBM primary cell cultures from the central U87 cells and the more clinically relevant invasive margin GIN-8 cells.

## Materials and Methods

### Reagents

The selective CB2 inverse-antagonist (AM630) was purchased from Tocris (Bristol, UK). All other reagents and materials for cell cultures were obtained from standard suppliers.

### Cellular Models

All cell lines (U87 and GIN-8) used in this project were of human origin, obtained from Dr Ruman Rahmam (University of Nottingham). U87 cells confirmed on Short Tandem Repeat (STR) genotyping, isolated from the core of a GBM tumor and sourced commercially, were used as a biological positive control for GBM cells. The Glioma INvasive margin (GIN-8) cell line, isolated from the tumor infiltrative edge, were derived in-house from surgeries at the Queen’s Medical Centre, Nottingham (comparable to their respective primary tissue on STR). Monolayer cells cultures were prepared as described previously[20, 21] Briefly, cells were plated into T75 cell culture flasks (Nunc, UK) until reaching confluency. Cells were trypsinazed and plated at a density of 25000 cells/ml in 6-multiwell plates in DMEM (Sigma, UK), supplemented with 10% fetal bovine serum (Sigma, UK), 5mM sodium pyruvate (Sigma, UK), 5mM L-Glutamine (Sigma, UK) and maintained in a humidified incubator at 37°C and 5% CO2. Cells grown in 6-well plates were treated with AM630 (5µM) or vehicle for 24h at 37°C. Cells then were harvested for microarray analysis.

### Microarray Analysis

Following cell treatment, the expression changes relative to control were assayed on a microarray chip with quadruplicate samples. Cells were lysed in Absolutely RNA Miniprep Kit lysis buffer and β-mercaptoethanol (Agilent Technologies, UK). RNA was then extracted, and quality assessed (RNA Integrity Number ≥ 8) using a Bioanalyzer (Agilent Technology). RNA expression levels were measured on Affymetrix Human Genome U133 plus 2.0 (GPL570) chip following library preparation and labelling by Nugen Ovation V2 and Nugen Encore as perthe supplier’s recommended procedure (https://www.selectscience.net/products/ovation-rna-seq-system-v2). Resulting expression data were pre-processed using RMA normalisation with the Bioconductor affy package[22].

### Expression analysis

The NCBI GEO hosts 145,000 samples of the Affymetrix Human Genome U133 Plus 2.0 Array on this platform, making it the most popular array chip. The relative expression levels of probes were collected for the GEO data and the given cell types. The ranks were scaled to lie between zero for the highest expression probe to unity for the lowest. The relative rank of each probe was defined as 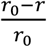 for *r*<*r*_0_and 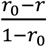, *r*<*r*_0_,where *r* and *r*_0_are the average probe ranks over the given cell type samples and the set of samples deposited on GEO respectively. Probes were then mapped to genes and in the case of degeneracy, the probe with the largest relative rank mapping to the gene. The resulting profile is the gene rank profile (GRP). This analysis is consistent with that presented by Hompoonsup et al[23].

The GRP corresponding to the untreated cell samples were queried against RNAseq data for a panel of cancer cell lines hosted by The Cancer Genome Atlas (TCGA) (https://cancergenome.nih.gov/). The panel consists of 7932 samples from 17 cancer types. Samples corresponding to a given cancer type were grouped and the cancer type gene expression levels assigned to the median of the group, resulting in 17 profiles of median expression. The gene expression levels were then ranked and compared to the ranked gene expression levels across the Genotype-Tissue Expression (GTEx) normal tissue RNAseq data (https://gtexportal.org/), to generate relative rank profiles similar to the microarray procedure discussed above. The Spearman correlation coefficient of the core and invasive margin cell GRP with the cancer type profiles were then generated.

Expression profiles contrasting the core and invasive boundary cell populations and for the cannabinoid inhibitor effects were based on the differences in group average probe expression levels, and ranked based on linear fit Z scores. The probes were mapped to genes with the maximal magnitude Z score slected in cases of alternative probes.

Profiles were compared with a Spearman rank correlation analysis and Fisher exact test across subsets of significantly regulated genes shared by the two profiles.

### Drug comparison

The AM630 profiles were queried against the Connectiity Map (CMAP)[24] data comprising the transcriptional profiles of 1,309 drug-like compounds using the SPIED platform[25]. The CMAP drugs were ranked according to the Fisher exact test for shared genes.

### Transcription factor co-expression profiles

Transcription factor co-expression profiles (TFCEP) were generated by collecting pairs of genes with the highest co-expression patterns across the 600,000 expression samples comprising SPIED[25]. Co-expression was measured by a Fisher exact test across samples for which both genes showed significant deviation from the sample series average. Each gene is assigned a profile consisting of the top 500 positively and negatively co-expressed genes. The transcription factor subset is obtained with reference to the gene ontology[26] assignment (GO:0003700).

### Pathway analysis

Pathway enrichment analysis was based on a variant of the Kolmogorov-Smirnoff (KS) statistic. The expression profile to be analysed was ordered based on the Z-score in the case of internally contrasted profiles, and the realtive rank in the case of profiles based on a comparison with external data. The KS measures the maximal displacement, *D*, from the null hypothesis distribution of a cumulative distribution of pathway genes on the ranked profile. The null distribution corresponds to the distribution of *D* for *N* random selections from an ordered set of *M*. Explicitly, for a cummulitive pathway gene count *C*_*i*_:

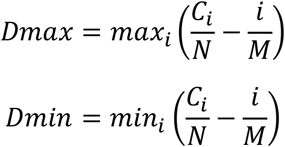

We observe that the distribution of *Dmax* + *Dmin* is normal with a standard deviation approximated by

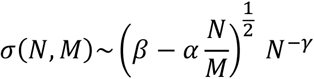

Where *α* = 0.3274679, *β* = 0.3327016 and *γ* = 0.491337. The reported statistic is the Z score 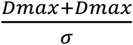.

## Results

### Characterising the GBM core and invasive margin cell populations

Cell functionality is encoded in the expressed genes and relative expression can be quantified by comparing the probe level ranks in the samples relative to their ranks across multiple samples profiled on the same array platform, to generate a GRP[23], see Methods. Comparing the GRP with similar profiles for multiple cancer cell lines from The Cancer Genome Atlas (TCGA) (https://cancergenome.nih.gov/) further strengthens our understanding of the basal transcriptomic profile of the core cell population, see Table 1. However, the invasive cell population has no correlation with any cell lines of this cancer panel, see Table 1. A pathway analysis of the two GRPs is shown in Table 2. Thus, it appears thatthe GBM infiltrative margin exhibits a unique transcriptomic profile relative to tumour core. The TCGA data is primary data from 206 patients with classical, mesenchymal and proneural subtypes of GBM.

**Table 1.** The rank expression profiles for the untreated cells compared to a panel of cancer cell types from the TCGA database. The rank expression profile for the untreated core cells was queried against a panel of 17 cancer cell lines from the TCGA database. The Spearman rank correlation coefficient is shown together with the Z score significance level. It is clear that the core cells correlate best with the GBM cell line from the TCGA database, shown left. The invasive GBM cell population, at right, has no positive correlation with any of the cancer cell types, including GBM.

| CORE |  |  | INVASIVE |  |  |
| --- | --- | --- | --- | --- | --- |
| CELL | r | z | CELL | r | z |
| GBM | 0.33 | 42.09 | SKCM | -0.05 | -6.72 |
| SKCM | 0.30 | 38.12 | UCEC | -0.06 | -7.02 |
| BRCA | 0.28 | 35.72 | GBM | -0.07 | -8.56 |
| LUSC | 0.27 | 34.09 | TGCT | -0.07 | -8.96 |
| HNSC | 0.26 | 32.79 | LIHC | -0.08 | -9.82 |
| BLCA | 0.26 | 32.64 | OV | -0.08 | -10.08 |
| TGCT | 0.25 | 32.62 | KIRP | -0.08 | -10.07 |
| OV | 0.25 | 31.90 | PAAD | -0.08 | -10.48 |
| CESC | 0.24 | 30.97 | READ | -0.08 | -10.48 |
| UCEC | 0.24 | 29.86 | HNSC | -0.09 | -11.21 |
| STAD | 0.23 | 29.94 | BLCA | -0.09 | -11.24 |
| PRAD | 0.23 | 29.57 | KICH | -0.09 | -11.65 |
| THCA | 0.21 | 26.94 | COAD | -0.09 | -11.72 |
| LUAD | 0.21 | 26.73 | CESC | -0.10 | -12.15 |
| COAD | 0.21 | 26.40 | PRAD | -0.10 | -12.61 |
| READ | 0.21 | 26.21 | THCA | -0.10 | -12.57 |
| KIRC | 0.21 | 26.40 | STAD | -0.12 | -14.59 |
| KICH | 0.21 | 25.85 | LUSC | -0.12 | -15.39 |
| PAAD | 0.20 | 25.56 | LUAD | -0.13 | -16.28 |
| KIRP | 0.19 | 23.91 | KIRC | -0.14 | -17.13 |
| LIHC | 0.17 | 20.84 | BRCA | -0.14 | -17.76 |

**Table 2.** Positively enriched pathways in the GPR profiles for the core and invasive margin cells. As expected, the core cells show a large positive enrichment with a series of cancer associated pathways, shown at left. In contrast, the invasive GPR enrichment scores are more modest, and reveal a distinctly unique pathway enrichment, relative to the core.

| CORE |  | INVASIVE |  |
| --- | --- | --- | --- |
| Z | PATHWAY | Z | PATHWAY |
| 13.98 | CHEK2 PCC NETWORK | 8.79 | ES WITH H3K27ME3 |
| 13.96 | TARGETS OF MIR192 AND MIR215 | 8.60 | MEF HCP WITH H3K27ME3 |
| 13.57 | MYCN TARGETS WITH E BOX | 7.59 | PRC2 TARGETS |
| 13.16 | BRCA1 PCC NETWORK | 7.36 | SUZ12 TARGETS |
| 12.58 | LUNG CANCER POOR SURVIVAL A6 | 7.08 | TRANSIENTLY UP BY 2ND EGF PULSE ONLY |
| 11.59 | NANOG TARGETS | 6.99 | EED TARGETS |
| 10.61 | ACINAR DEVELOPMENT LATE 2 | 6.94 | MCV6 HCP WITH H3K27ME3 |
| 10.61 | CYCLING GENES | 6.34 | NPC HCP WITH H3K4ME2 AND H3K27ME3 |
| 10.45 | RB1 TARGETS SENESCENT | 5.74 | BRAIN HCP WITH H3K27ME3 |
| 10.25 | TARGETS OF SMAD2 OR SMAD3 | 5.63 | NPC HCP WITH H3K27ME3 |
| 9.95 | LIVER CANCER | 5.58 | EBNA1 ANTICORRELATED |
| 9.95 | BRCA2 PCC NETWORK | 5.49 | TP63 TARGETS |
| 9.92 | BOUND BY E2F4 UNSTIMULATED | 5.16 | EGF RESPONSE 480 HELA |
| 9.60 | MIR21 TARGETS | 5.16 | DILATED CARDIOMYOPATHY |
| 9.33 | RB1 TARGETS GROWING | 4.87 | GASTRIC CANCER CHEMOSENSITIVITY |
| 9.10 | RB1 TARGETS CONFLUENT | 4.84 | BREAST CANCER 16P13 AMPLICON |
| 8.99 | EMBRYONIC STEM CELL CORE | 4.79 | FOCAL ADHESION |
| 8.91 | SOX2 TARGETS | 4.31 | ES ICP WITH H3K27ME3 |
| 8.88 | TARGETS OF MIR34B AND MIR34C | 4.21 | MBD TARGETS |
| 8.69 | HYPOXIA NOT VIA KDM3A | 4.20 | NFAT 3PATHWAY |

It is clear that the core GBM cells express high levels of the CB1 receptor (CNR1) but not the CB2 receptor (CNR2), see Table 3. This led us to investigate alternative targets for AM630 activity. A comprehensive investigation of CNR2 ligand profiling[27] delimited the off-target activity of AM630. In total 11 non-CB2 targets were identified: CNR1, TRPA1, A3 receptor (ADORA3), GABA-gated Cl-channel (GABRA1), FP (PTGFR), 5-HT2A/ B receptors (HTR2A andHTR2B), KOP (OPRK1), PPARG, COX2 (PTGS2). Of these CNR1 shows the highestest expression within the core cell population. Whereas in the invasive margin cells, neither of the CB receptors are expressed. The only putative target for AM630 engagement with the invasive margin cells are the 5-HT serotonin receptors (HTR2A and HTR2B) and the transient receptor potential channel TRPA1. The serotonin receptors have moderate expression in both cell populations and we expect any shared response to AM630 to be through inhibition of these receptors. By virtue of the high expression of the CB1 receptor in the core cells it is reasonable to expect thet these cells will be more responsive to AM630 than the invasive margin cells. This exemplifies the caution needed when predicating preclinical drug selection and dosing, upon data generated from the GBM core. Our data supports the notion that drug selection/dosing should be informed by the clinically-relevant infiltrative margin of GBM.

**Table 3.**
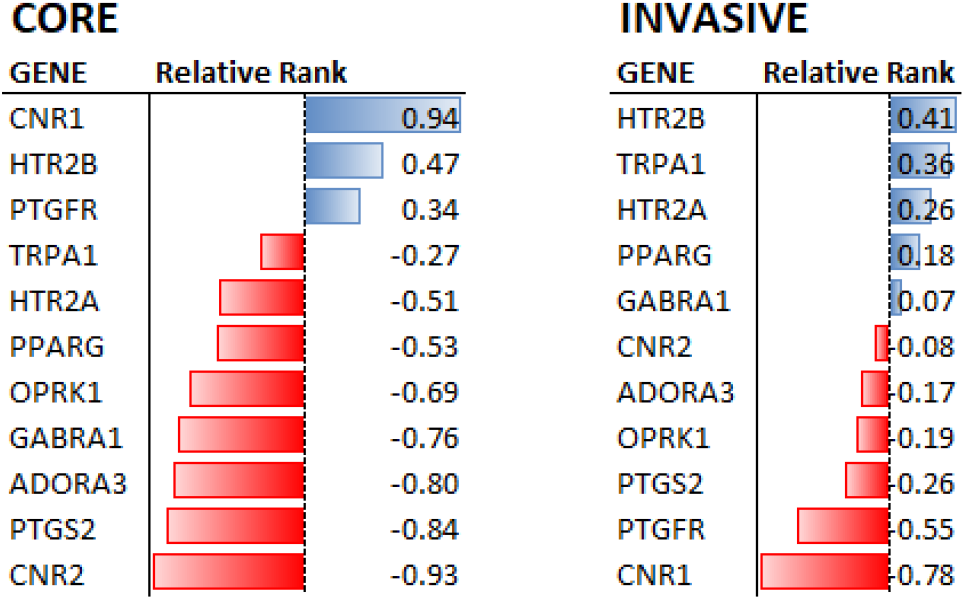
The relative expression levels of candidate targets for AM630 in the core and invasive cell populations. Genes that are reported as off-target for AM630 are expressed at different levels in the two cell populations with the CB1 receptor (CNR1) showing the highest expression in the core cells indicating that the dominant activity elicited by AM630 in this population will most likely be mediated through this receptor, shown at left. In contrast the invasive margin cells do not have conspicuously high CNR1 expression, shown at right. Only the 5-HT serotonin receptor HTR2B is upregulated in both populations, suggesting that a shared activity of AM630 might involve serotonin receptor antagonism.

### Differential expression induced by AM630

The CB antagonist AM630 drives a substantial transcriptional response in the core cell population. The transcriptional response was symmetrical and substantial with 1672 and 2424 two-fold up and downregulated genes, see Figure 1 and Table S1. This is in contrast to the invasive margin cells where there are 815 and 660 two-fold up and down regulated genes.

**Figure 1.**
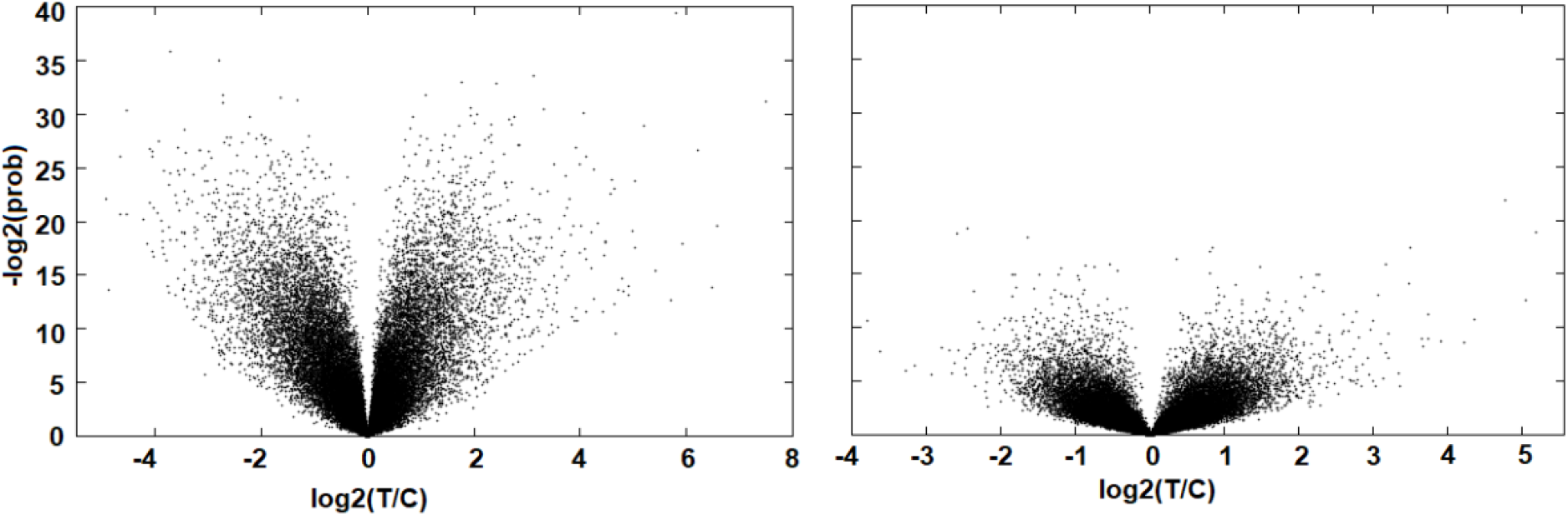
Volcano plot of genes significantly perturbed by AM630. The response is substantial in the core cells (left) but relatively dampened in the invasive margin cells (right).

The muted transcriptional response to AM630 in the invasive margin population has only a moderate correlation with the core cell response. Comparing probes with significant expression differences (at the three standard deviation from the null level) between treatment and control in the two cell populations we see a slight correlation in the AM630 response, see Figure 2. This observation might speak to the similar expression level of serotonon receptors in the two cell populations.

**Figure 2.**
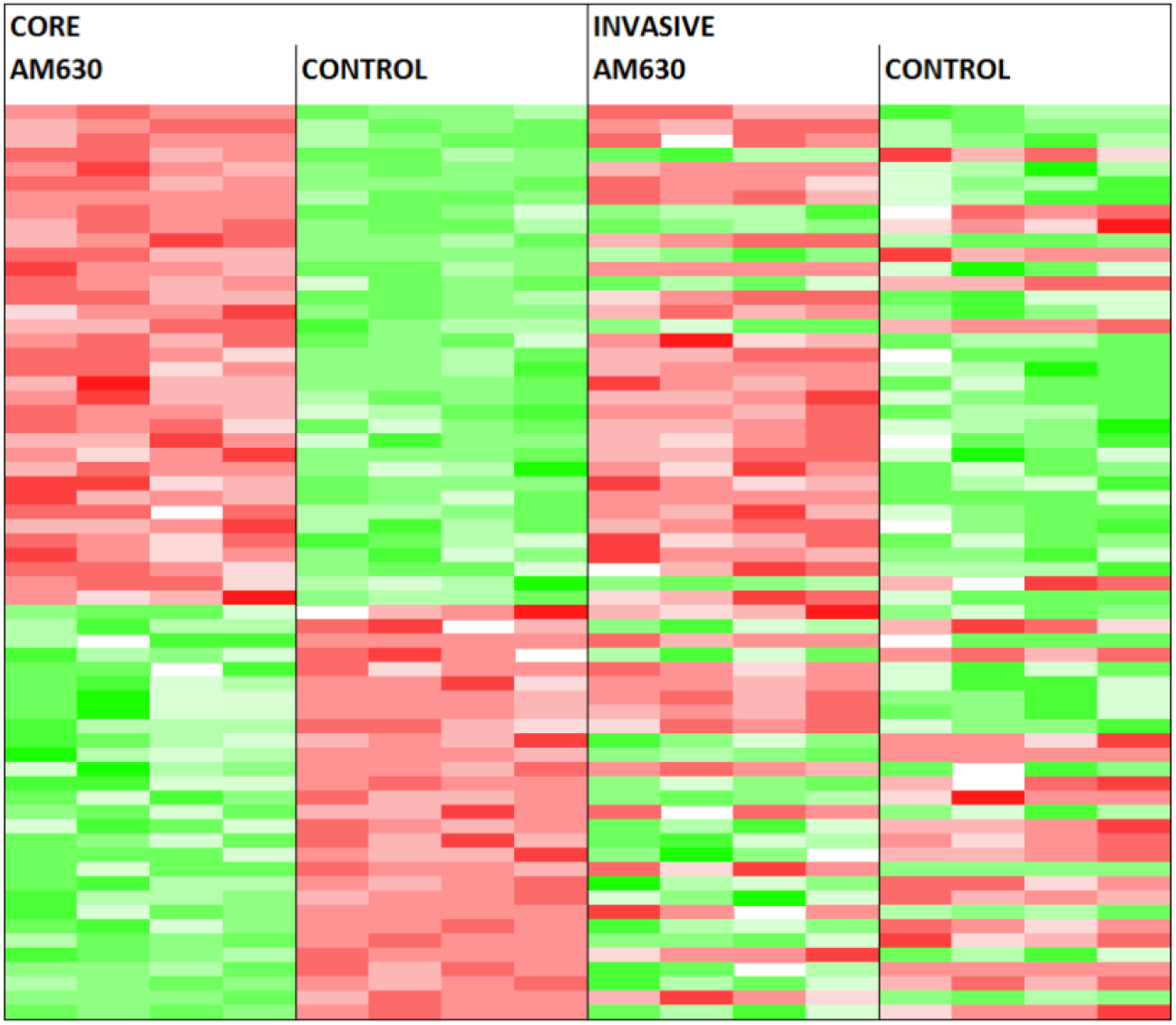
The probes that are significantly perturbed by AM630 in both the core and invasive margin cells. Probe levels are shown for which both the AM630 core and invasive cell expression change is above three standard deviations from the null. The total number of probes is 64 and there is a significant correlation between the responses in both cell types, with a Fisher exact test score of p < 0.004 (UU 28, UD 7, DU 13, DD 16, where U = up regulated and D = down regulated).

Pathway enrichment of the genes perturbed in the core population shows a clear up regulation of immune response genes and the down regulation of cell cycle and metastatic pathways, see Table 4. In contrast the weaker respose in the invasive margin is associated with more modest pathway enrichment, see Table 5. However, of note is the down regulation of tumour invasivenes genes and the MYC oncogene signature indicating an anti-neoplastic AM630 activity in this context.

**Table 4.**
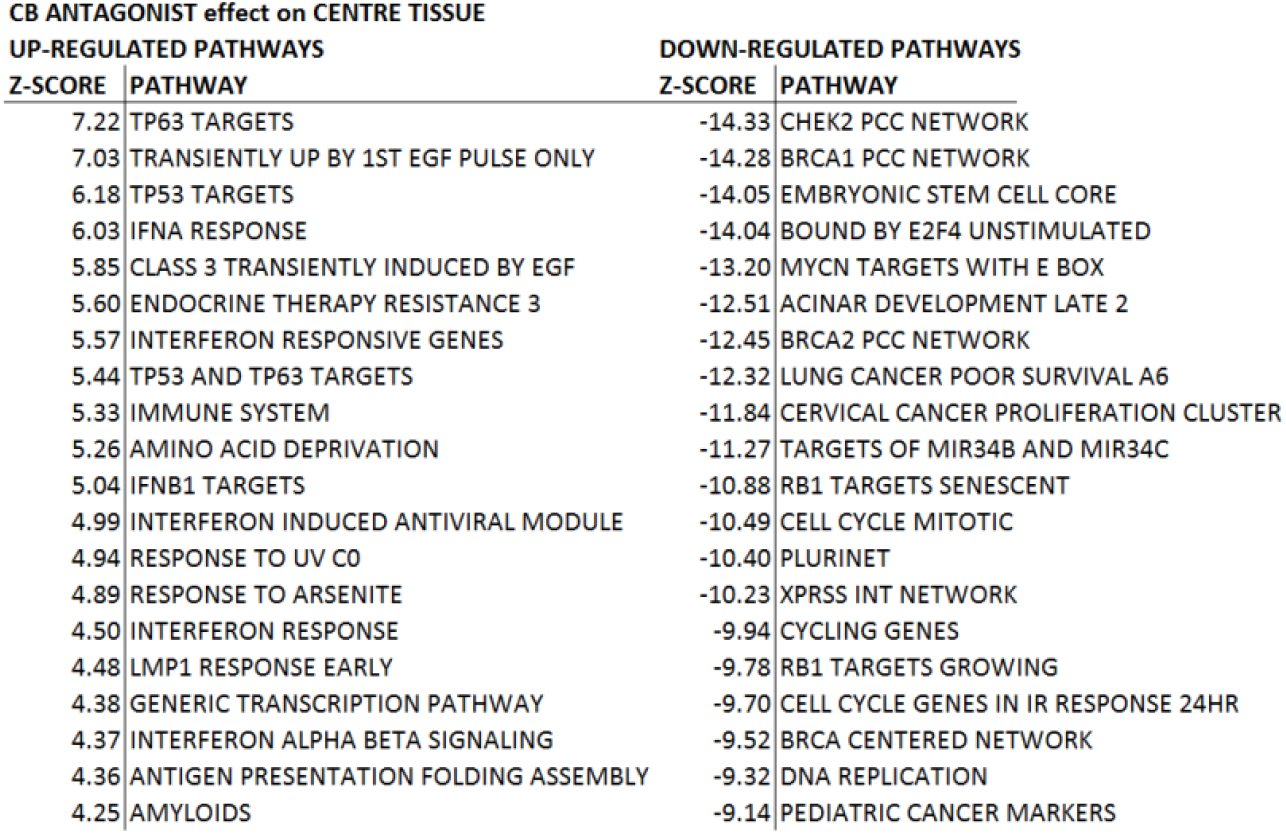
The positive and negative enrichment of pathways in the AM630 transcriptional response of the core cell population. AM630 up regulates an immune response together with TP53 and interferon and a down regulation of cell cycle and cancer associated pathways.

**Table 5.**
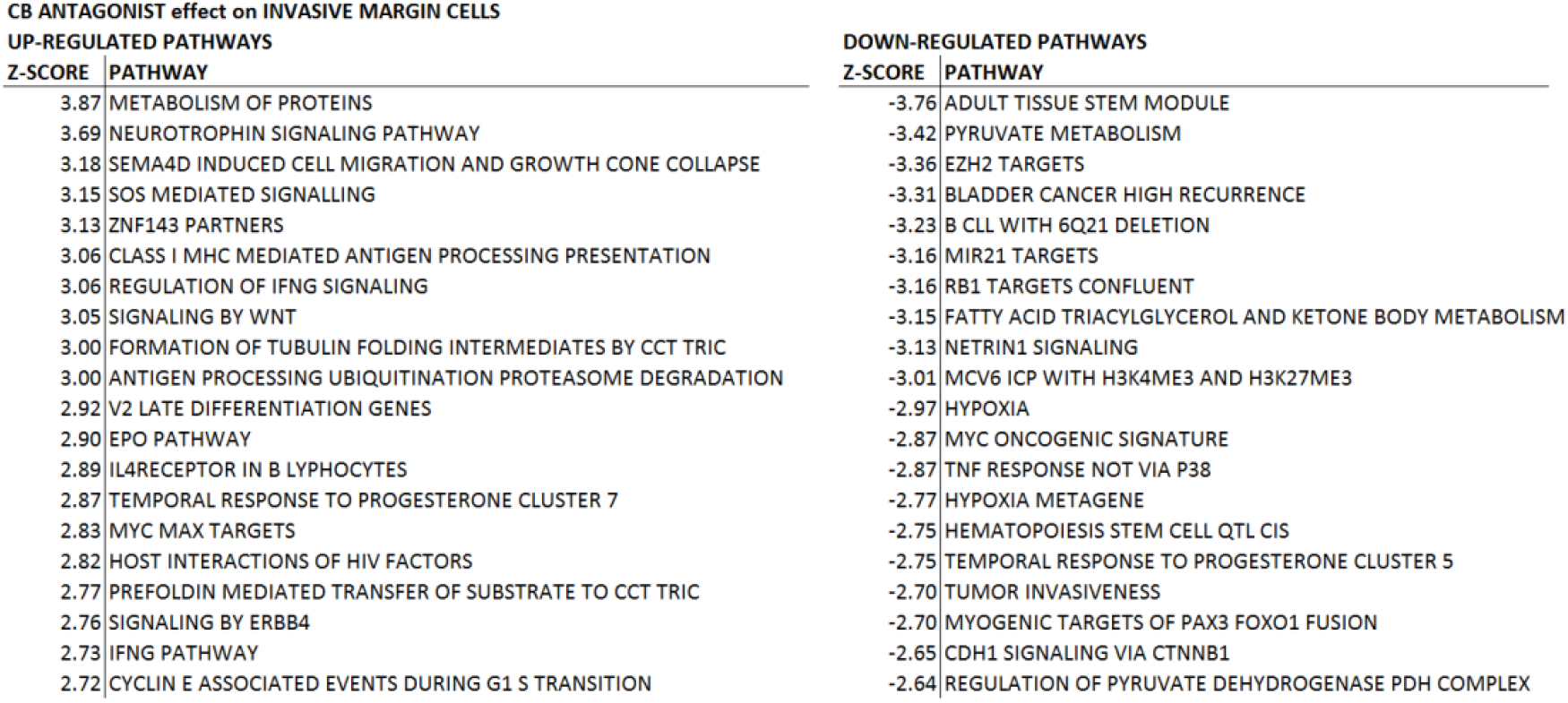
The positive and negative enrichment of pathways in the AM630 transcriptional response of the invasive cell population. The invasive margin response is associated with the enrichment of distinct pathways relative to the core cell response. Of note are the down regulation of the tumour invasiveness and MYC pathways, indicating a possible anti-neoplastic activity.

| CB ANTAGONIST effect on INVASIVE MARGIN CELLS |  | DOWN-REGULATED PATHWAYS |  |
| --- | --- | --- | --- |
| UP-REGULATED PATHWAYS |  |  |  |
| Z-SCORE | PATHWAY | Z-SCORE | PATHWAY |
| 3.87 | METABOLISM OF PROTEINS | -3.76 | ADULT TISSUE STEM MODULE |
| 3.69 | NEUROTROPHIN SIGNALING PATHWAY | -3.42 | PYRUVATE METABOLISM |
| 3.18 | SEMA4D INDUCED CELL MIGRATION AND GROWTH CONE COLLAPSE | -3.36 | EZH2 TARGETS |
| 3.15 | SOS MEDIATED SIGNALLING | -3.31 | BLADDER CANCER HIGH RECURRENCE |
| 3.13 | ZNF143 PARTNERS | -3.23 | B CLL WITH 6Q21 DELETION |
| 3.06 | CLASS I MHC MEDIATED ANTIGEN PROCESSING PRESENTATION | -3.16 | MIR21 TARGETS |
| 3.06 | REGULATION OF IFNG SIGNALING | -3.16 | RB1 TARGETS CONFLUENT |
| 3.05 | SIGNALING BY WNT | -3.15 | FATTY ACID TRIACYLGLYCEROL AND KETONE BODY METABOLISM |
| 3.00 | FORMATION OF TUBULIN FOLDING INTERMEDIATES BY CCT TRIC | -3.13 | NETRIN1 SIGNALING |
| 3.00 | ANTIGEN PROCESSING UBIQUITINATION PROTEASOME DEGRADATION | -3.01 | MCV6 ICP WITH H3K4ME3 AND H3K27ME3 |
| 2.92 | V2 LATE DIFFERENTIATION GENES | -2.97 | HYPOXIA |
| 2.90 | EPO PATHWAY | -2.87 | MYC ONCOGENIC SIGNATURE |
| 2.89 | IL4RECEPTOR IN B LYMPHOCYTES | -2.87 | TNF RESPONSE NOT VIA P38 |
| 2.87 | TEMPORAL RESPONSE TO PROGESTERONE CLUSTER 7 | -2.77 | HYPOXIA METAGENE |
| 2.83 | MYC MAX TARGETS | -2.75 | HEMATOPOIESIS STEM CELL QTL CIS |
| 2.82 | HOST INTERACTIONS OF HIV FACTORS | -2.75 | TEMPORAL RESPONSE TO PROGESTERONE CLUSTER 5 |
| 2.77 | PREFOLDIN MEDIATED TRANSFER OF SUBSTRATE TO CCT TRIC | -2.70 | TUMOR INVASIVENESS |
| 2.76 | SIGNALING BY ERBB4 | -2.70 | MYOGENIC TARGETS OF PAX3 FOXO1 FUSION |
| 2.73 | IFNG PATHWAY | -2.65 | CDH1 SIGNALING VIA CTNNB1 |
| 2.72 | CYCLIN E ASSOCIATED EVENTS DURING G1 S TRANSITION | -2.64 | REGULATION OF PYRUVATE DEHYDROGENASE PDH COMPLEX |

Another insight into the activity of AM630 can be gained by a direct correlation analysis with the transcriptional activity of other compounds. To this end we performed a search of the Connectivity Map (CMAP)[24] database of 1,300 drug-like compounds profiled in cancer cell lines through the SPIED platform (www.spied.org.uk)[25]. As expected, the highly correlated profiles correspond to anti-proliferative agents with some having reported inhibitory effects against GBM, see Table 6. Conspicuos in the list of CMAP profiles correlating with the AM630 core profile are the six antipsychotic serotonin receptor inhibitors: thioridazine, fluphenazine, prochlorperazine, perphenazine, nortriptyline, metergoline. Interestingly, antipsychotics have been suggested as repurposing candidates for GBM[28, 29]. Specifically, perphenazine and prochlorperazine have sub-micromolar cytotoxixity against the U87MG GBM cell line[30] with thioridazine and fluphenazine having reported anti GBM8401 and U87MG GBM cell activity[31]. Metergoline was reported as a GBM stem cell proliferation inhibitor in a high-throughput screen[32]. Another class of drugs scoring highly against the AM630 profile are the HDAC inhibitors: scriptaid, vorinostat, trichostatin A. HDAC inhibition has also been proposed as an intervention in GBM[33] with vorinostat reaching phase II clinical trails for GBM[34] and scriptaid inducing glioma cell apoptosis[35]. The top correlating drug, prostaglandin J2, is a PPARg agonists and has shown anti glioma activity. The PI3 kinase inhibitor LY-294002 has been reported to inhibit the growth of malignant glioma cells[36].

**Table 6.**
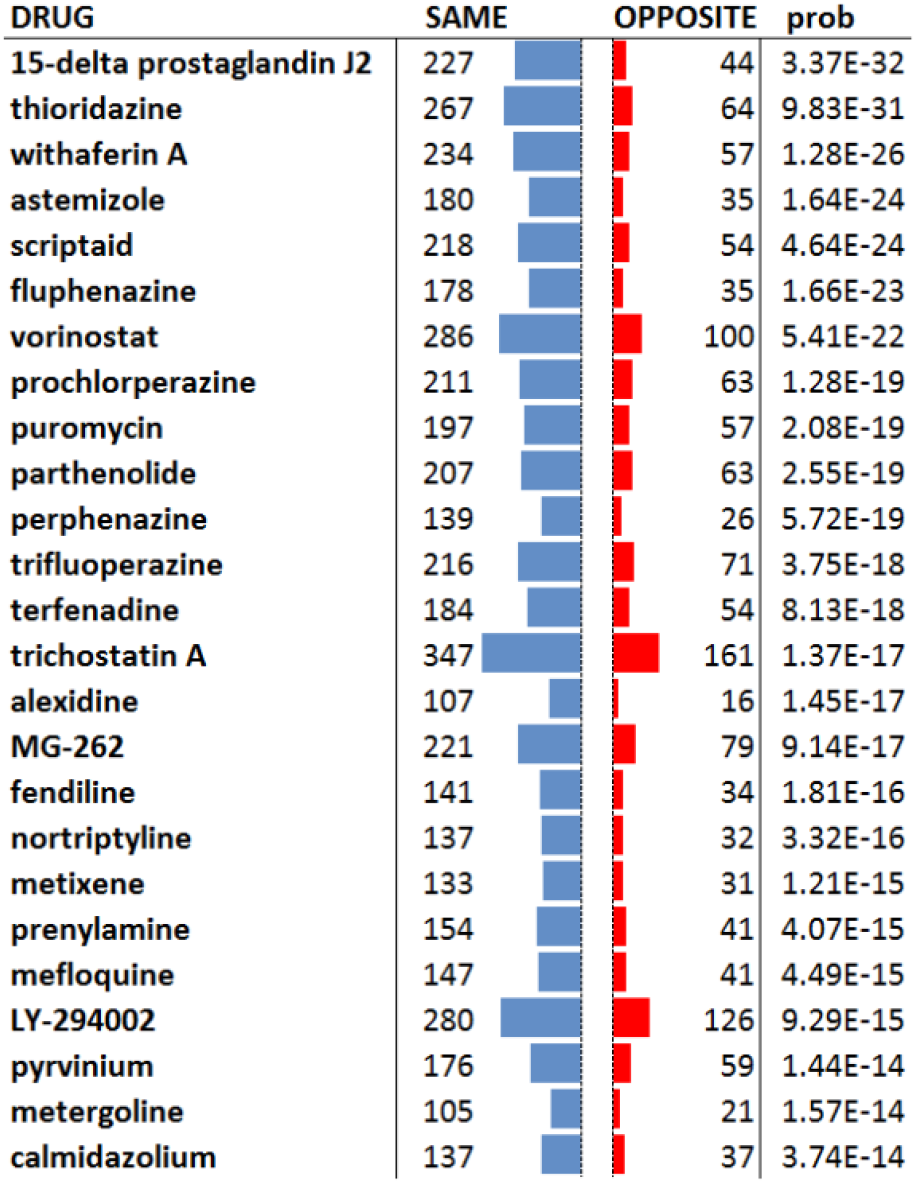
Drug-like compound expression profiles that correlate with AM630 driven expression changes in the core cell population. At the five standard deviations from the null level there are 343 up and 348 down regulated genes. There is a substantial correlation with established antineoplastics and compounds with specific activity against GBM. Conspicuous amongst the correlating drugs are the six antipsychotic serotonin receptor antagonists: thioridazine, fluphenazine, prochlorperazine, perphenazine, nortriptyline, metergoline.

| DRUG | SAME | OPPOSITE | prob |
| --- | --- | --- | --- |
| 15-delta prostaglandin J2 | 227 | 44 | 3.37E-32 |
| thioridazine | 267 | 64 | 9.83E-31 |
| withaferin A | 234 | 57 | 1.28E-26 |
| astemizole | 180 | 35 | 1.64E-24 |
| scriptaid | 218 | 54 | 4.64E-24 |
| fluphenazine | 178 | 35 | 1.66E-23 |
| vorinostat | 286 | 100 | 5.41E-22 |
| prochlorperazine | 211 | 63 | 1.28E-19 |
| puromycin | 197 | 57 | 2.08E-19 |
| parthenolide | 207 | 63 | 2.55E-19 |
| perphenazine | 139 | 26 | 5.72E-19 |
| trifluoperazine | 216 | 71 | 3.75E-18 |
| terfenadine | 184 | 54 | 8.13E-18 |
| trichostatin A | 347 | 161 | 1.37E-17 |
| alexidine | 107 | 16 | 1.45E-17 |
| MG-262 | 221 | 79 | 9.14E-17 |
| fendiline | 141 | 34 | 1.81E-16 |
| nortriptyline | 137 | 32 | 3.32E-16 |
| metixene | 133 | 31 | 1.21E-15 |
| prenylamine | 154 | 41 | 4.07E-15 |
| mefloquine | 147 | 41 | 4.49E-15 |
| LY-294002 | 280 | 126 | 9.29E-15 |
| pyrvinium | 176 | 59 | 1.44E-14 |
| metergoline | 105 | 21 | 1.57E-14 |
| calmidazolium | 137 | 37 | 3.74E-14 |

In contrast a CMAP analysis of the invasive AM630 margin profile does not return many significantly correlating drugs. Noting the moderate correlation in the core and invasive margin responses we reasoned that a combined profile populated with genes with a significant combined expression change may capture the shared activity in the two cellular contexts. To this end we generated a combined profile constituting genes with combined Z scores of above five standard deviations from the null using Stouffer’s method[37], see Table S2. Here, there is a significant overlap in the core cell response result, noteably the serotonin receptor antagonsits are still significantly correlated.

Large scale transcriptional data enables the discovery of functional associations between genes[38], where gene pairs with correlated expression changes across multiple experiments are likely to be involved in similar biological functions. This enabled us to pinpoint transcription factors whose regulation may recapitulate the inhibitory activity of AM630, see Methods. Table 7 shows the top positively and negatively correlating transcription factors. This analysis suggests that AM630 activity might be recapitulated by the up regulation of DDIT3 in GBM. This is of interest, because DDIT3 expression leads to the modulation of NAG-1, resulting in GBM cell apoptosis[39]. Similarly, another TF involved in apoptosis through the ER stress pathway, CREBRF, is positively correlated with AM630 activity[40]. It would be of interest to investigate the therapeutic potential of inhibiting the negatively correlated TFs. For example FOXM1 induces resistance to radiotherapy by modulating the activity of SOX-2[41]. Not surprisingly we see a correlation with the down regulation of multiple E2F TFs as these are required for cell cycle progression[42] and associated with GBM malignancy progression[43]. Inhibition of the Fanconi Anaemia pathway gene FANCD2 has been reported to sensitise gliomas to chemotherapeutic intervention[44]. GBM is also strongly associated with the expression of TFDP1[45].

**Table 7.**
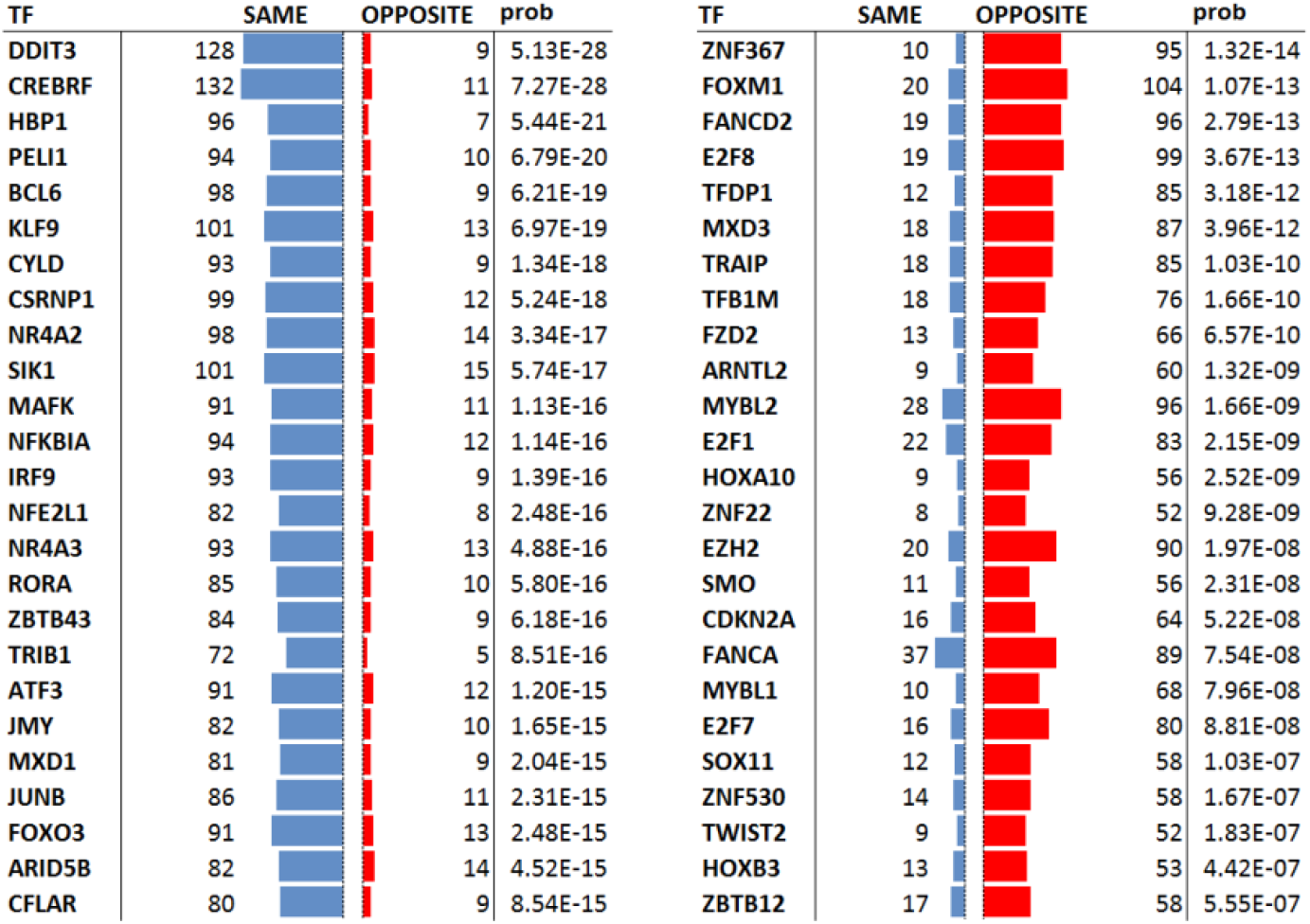
Transcription factors predicted to recapitulate the inhibitory activity of AM630. The AM630 transcription profile in the core cells restricted to 343 up and 348 down regulated genes passing the five standard deviations away from the null significance level, were queried against TFCEPs (see Methods). The positively correlating TFs are shown on the left and negatively correlating TFs on the right. Up/down regulation of the positively/negatively correlating TFs are hypothesised to recapitulate the inhibitory activity of AM630.

## Discussion

Despite multiple clinical trials and research efforts, GBM continues being the most aggressive of all forms of cancer with very limited clinical options to stop progression and dissemination of the tumor. GBM constitutes a complex of interacting cell types with a core population responding to treatments and an invasive margin population that has proved refractory to intervention[46]. Unfortunately, all efforts to cure this type of cancer have failed to significantly extend median survival times[4].

Different signalling pathways associated with the brain endocannabinoid machinery are being investigated as potential therapeutic targets[47], with recent evidence suggests that blocking cannabinoid machinery mediates antitumour effects via the inactivation of traditional cannabinoid receptors (CB1 or CB2)[13, 14]. This is a research field in development, with limited results, as the molecular mechanisms of this anti-tumoral effect downstream of the cannabinoid receptors activation are unknown[48]. In this context, we have previously reported on the effectiveness of the CB2 receptor inverse agonist AM630 in blocking core GBM cell proliferation[18]. However, AM630 appears to be a less potent inhibitor of the invasive margin cell population. To uncover the biological mechanisms underlying AM630 activity we performed a gene expression analysis of treated core, U87 cells, and invasive margin, GIN-8 cells, of human GBM.

A relative expression analysis of the untreated cells highlights the differences between the core and invasive margin populations. The core cells show a clear correlation with published GBM expression data whereas the invasive margin population shows no correlation with any of the TCGA cell lines. This is in agreement with recent evidence that reveals an extensive degree of intra-tumour heterogeneity in GBM resulting from a clonal evolution process producing a completely different genetic pattern in the core compared with the invasive margin[46]. This different molecular signature is critical for the selection potential pharmacotherapies for clinical application. The margin cells, which ultimately result in the inevitable recurrence of GBM, escape from traditional treatments[46]. Residual cells at the tumor invasive margin are responsible for the 85% of GBMs that relapse locally following resection plus radiotherapy and temozolomide[49].

As regards the potential activity of AM630, we found low primary target, the CB2 receptor, expression in either population. This led us to consider potential off-target effects for AM630 activity. Of the off-targets the CB1 receptor is highly expressed in the core and the serotonin receptor (HTR2B) expressed at moderate levels in both populations. This observation is in agreement with the relatively high potency of AM630 in the core population. We hypothesise that AM630 elicits a substantial response in the core cells through antagonism of the CB1 receptor and any activity shared by the drug across the core and invasive margin cells may be driven by serotonin receptor antagonism.

Analizing the pathways involved, we found that in the core of the tumor, AM630 is associated with the down regulatiuon of cell cycle and cancer associated pathways. In contrast, AM630 up-regulates an immune response together with the TP53 and interferon pathways. The TP53 gene encodes a tumor suppressor protein P53, that plays a critical role in tumor suppression by orchestrating a wide variety of cellular responses inducing tumoral cell death[50]. We can speculate that AM630 effects in the core of the tumor are a restoration of P53 function, inducing a change in the cytokine network, mainly via IFN signalling pathways. Consecuently, the above restorative and pro-inflammatory AM630 actions, will induce cell cycle arrest and cell death of GBM. Our findings about the beneficial effects of harnessing the IFN signallind pathway, are supported by recent evidence that identify an IFN-β-associated gene signature as a marker for the prediction of overall survival among glioblastoma patients[51]. In tandem with the pathway analysis we performed an investigation of the transcription factors predicted to recapitulate the inhibitory activity of AM630. Of note is the positive correlation with the DNA damage induced TF, DDIT3, because DDIT3 expression leads to the modulation of NAG-1, and the whole process results in GBM cell apoptosis[39]. Similarly, CREBRF, is a TF involved in inducing cell apoptosis through the ER stress pathway[40]. TFs whose down regulation tends to drive gene expression in the direction of the AM630 response may emerge as therapeutic targets for inhibition. This is boltered by the observation that FOXM1 that induces resistance to radiotherapy by modulating the activity of SOX-2[41], also the TFs FANCD2 and E2F8 have expression levels strongly associated with GBM malignancy progression[43] as has TFDP1[45].

Investigating the drug driven transcriptional profiles from the CMAP database we found that anti-neoplastic agents have high levels of correlation with the core cell response profile, with some drugs reported to have anti GBM activities. Of particular interest was the high degree of correlation with serotonin receptor antagonists which may speak to the serotonin receptor HTR2B being the sole off-target of AM630 with significant expression in both the core and invasive margin cell populations. However, the relatively weak AM630 response in the invasive margin population has only a moderate correlation with the core cell response. A pathway analysis of the invasive margin response reveals the down regulation of pathways in associated with cancer, but a CMAP analysis does not return significantly drugs. However, a transcriptional profile constituting genes that are regulated in common in the two cell populations largely recapitulates the results in the core cell population. Specifically, the serotonin receptor antagonists correlate with the combined profile.

The correlation with serotonin receptor antagonists is of interest as the 5-HT serotonin receptors are highly expressed in different types of cancers, including GBM and modulate mitogenic signalling and impact tumour cell viability[52]. In this context, serotonin receptor antagonism has been hypotheises as a therapeutic intervention in GBM, with perphenazine and prochlorperazine showing inhibition of the U87MG GBM cell line[30], thioridazine and fluphenazine inhibiting both GBM8401 and U87MG GBM cell activity[31]. Metergoline was reported as a GBM stem cell proliferation inhibitor in a high-throughput screen[32]. The Inhibition of SMPD1, a gene that regulates ‘ceramide sphingosine-1-phosphate rheostat’ and drives tumor growth and immune escape in different types of cancer[53], through inhibition of epidermal growth factor receptor (EGFR) signalling and via activation of lysosomal stress has been proposed as the potential anti-tumoral effects of serotonin receptor inhibition, through fluoxidine, in GBM[54]. In preclinical studies, the effect of 5-HTR2B antagonists on angiogenesis was associated with decreased tumour microvessel density[55]. The substantial involvement of serotonin receptors, especially 5HTR2B, in different types of cancers[56], support further studies as a potential treatment target for both the core and invasive margin of GBM. Interestingly in our study, the HTR2B paralog gene HTR2A is substantially expressed in the boundary of the tumor.

Given the heterogeneity of glioblastoma, further studies are required to elucidated the molecular mechanisms of the observed AM630 anti-tumoral actions and if can potentially be used in the future as an addition to current therapy.

## Supporting information

SuppementaryTable S1 &S2

## Authors Contribution

All Authors have made a substantial contribution to this project, and have approved the submitted version of the manuscript.

## References

1. Garcia-Arencibia, M., E. Molina-Holgado, and F. Molina-Holgado, Effect of endocannabinoid signalling on cell fate: life, death, differentiation and proliferation of brain cells. Br J Pharmacol, 2019. 176(10): p. 1361–1369.

2. Fares, J., et al., The Network of Cytokines in Brain Metastases. Cancers (Basel), 2021. 13(1).

3. Smith, S.J., et al., Overall Survival in Malignant Glioma Is Significantly Prolonged by Neurosurgical Delivery of Etoposide and Temozolomide from a Thermo-Responsive Biodegradable Paste. Clin Cancer Res, 2019. 25(16): p. 5094–5106.

4. Aldape, K., et al., Challenges to curing primary brain tumours. Nat Rev Clin Oncol, 2019. 16(8): p. 509–520.

5. Batash, R., et al., Glioblastoma Multiforme, Diagnosis and Treatment; Recent Literature Review. Curr Med Chem, 2017. 24(27): p. 3002–3009.

6. Kano, M., et al., Endocannabinoid-mediated control of synaptic transmission. Physiol Rev, 2009. 89(1): p. 309–80.

7. Mechoulam, R. and L.A. Parker, The endocannabinoid system and the brain. Annu Rev Psychol, 2013. 64: p. 21–47.

8. Molina-Holgado, E. and F. Molina-Holgado, Mending the broken brain: neuroimmune interactions in neurogenesis. J Neurochem, 2010. 114(5): p. 1277–90.

9. Cahoy, J.D., et al., A transcriptome database for astrocytes, neurons, and oligodendrocytes: a new resource for understanding brain development and function. J Neurosci, 2008. 28(1): p. 264–78.

10. Irving, A., et al., Cannabinoid Receptor-Related Orphan G Protein-Coupled Receptors. Adv Pharmacol, 2017. 80: p. 223–247.

11. Guzman, M., Effects on cell viability. Handb Exp Pharmacol, 2005(168): p. 627–42.

12. Maccarrone, M., et al., Programming of neural cells by (endo)cannabinoids: from physiological rules to emerging therapies. Nat Rev Neurosci, 2014. 15(12): p. 786–801.

13. Malfitano, A.M., et al., The CB1 receptor antagonist rimonabant controls cell viability and ascitic tumour growth in mice. Pharmacol Res, 2012. 65(3): p. 365–71.

14. Proto, M.C., et al., Interaction of endocannabinoid system and steroid hormones in the control of colon cancer cell growth. J Cell Physiol, 2012. 227(1): p. 250–8.

15. Alberich Jorda, M., et al., The peripheral cannabinoid receptor Cb2, frequently expressed on AML blasts, either induces a neutrophilic differentiation block or confers abnormal migration properties in a ligand-dependent manner. Blood, 2004. 104(2): p. 526–34.

16. Wang, J., et al., Cannabinoid receptor 2 as a novel target for promotion of renal cell carcinoma prognosis and progression. J Cancer Res Clin Oncol, 2018. 144(1): p. 39–52.

17. Perez-Gomez, E., et al., Role of cannabinoid receptor CB2 in HER2 pro-oncogenic signaling in breast cancer. J Natl Cancer Inst, 2015. 107(6): p. djv077.

18. Nikoloudakou, A., Molina-Holgado, F, AM630, a CB2 cannabinoid receptor antagonist, inhibits proliferation of human U87 glioblastoma cells by targeting the mitochondrial unfolded protein response. Proceedings of the British Pharmacological Society, 2017. 17.

19. Sottoriva, A., et al., Intratumor heterogeneity in human glioblastoma reflects cancer evolutionary dynamics. Proc Natl Acad Sci U S A, 2013. 110(10): p. 4009–14.

20. Smith, S.J., et al., Metabolism-based isolation of invasive glioblastoma cells with specific gene signatures and tumorigenic potential. Neurooncol Adv, 2020. 2(1): p. vdaa087.

21. Smith, S.J., et al., Recapitulation of tumor heterogeneity and molecular signatures in a 3D brain cancer model with decreased sensitivity to histone deacetylase inhibition. PLoS One, 2012. 7(12): p. e52335.

22. Gautier, L., et al., affy--analysis of Affymetrix GeneChip data at the probe level. Bioinformatics, 2004. 20(3): p. 307–15.

23. Hompoonsup, S., et al., No transcriptional evidence for active Nav channels in two classes of cancer cell. Channels (Austin), 2019. 13(1): p. 311–320.

24. Lamb, J., et al., The Connectivity Map: using gene-expression signatures to connect small molecules, genes, and disease. Science, 2006. 313(5795): p. 1929–35.

25. Williams, G., SPIEDw: a searchable platform-independent expression database web tool. BMC Genomics, 2013. 14: p. 765.

26. Carbon, S., et al., AmiGO: online access to ontology and annotation data. Bioinformatics, 2009. 25(2): p. 288–9.

27. Soethoudt, M., et al., Protocol to Study beta-Arrestin Recruitment by CB1 and CB2 Cannabinoid Receptors. Methods Mol Biol, 2016. 1412: p. 103–11.

28. Lee, J.K., D.H. Nam, and J. Lee, Repurposing antipsychotics as glioblastoma therapeutics: Potentials and challenges. Oncol Lett, 2016. 11(2): p. 1281–1286.

29. Kamarudin, M.N.A. and I. Parhar, Emerging therapeutic potential of anti-psychotic drugs in the management of human glioma: A comprehensive review. Oncotarget, 2019. 10(39): p. 3952–3977.

30. Otreba, M. and E. Buszman, Perphenazine and prochlorperazine induce concentration-dependent loss in human glioblastoma cells viability. Pharmazie, 2018. 73(1): p. 19–21.

31. Cheng, H.W., et al., Identification of thioridazine, an antipsychotic drug, as an antiglioblastoma and anticancer stem cell agent using public gene expression data. Cell Death Dis, 2015. 6: p. e1753.

32. Visnyei, K., et al., A molecular screening approach to identify and characterize inhibitors of glioblastoma stem cells. Mol Cancer Ther, 2011. 10(10): p. 1818–28.

33. Chen, R., et al., The application of histone deacetylases inhibitors in glioblastoma. J Exp Clin Cancer Res, 2020. 39(1): p. 138.

34. Galanis, E., et al., Phase I/II trial of vorinostat combined with temozolomide and radiation therapy for newly diagnosed glioblastoma: results of Alliance N0874/ABTC 02. Neuro Oncol, 2018. 20(4): p. 546–556.

35. Sharma, V., et al., HDAC inhibitor, scriptaid, induces glioma cell apoptosis through JNK activation and inhibits telomerase activity. J Cell Mol Med, 2010. 14(8): p. 2151–61.

36. Shingu, T., et al., Growth inhibition of human malignant glioma cells induced by the PI3-K-specific inhibitor. J Neurosurg, 2003. 98(1): p. 154–61.

37. Stouffer, S.A., Suchman, E. A., DeVinney, L. C., Star, S. A., & Williams, R. M., Jr., The American Soldier: Adjustment During Army Life (Vol. 1). 1949: Princeton, NJ: Princeton University Press.

38. Williams, G., Database of Gene Co-Regulation (dGCR): A Web Tool for Analysing Patterns of Gene Co-regulation across Publicly Available Expression Data. J Genomics, 2015. 3: p. 29–35.

39. Tsai, S.F., et al., Isochaihulactone-induced DDIT3 causes ER stress-PERK independent apoptosis in glioblastoma multiforme cells. Oncotarget, 2017. 8(3): p. 4051–4061.

40. Yang, Y., et al., Luman recruiting factor regulates endoplasmic reticulum stress in mouse ovarian granulosa cell apoptosis. Theriogenology, 2013. 79(4): p. 633-9 e1-3.

41. Lee, Y., et al., FoxM1 Promotes Stemness and Radio-Resistance of Glioblastoma by Regulating the Master Stem Cell Regulator Sox2. PLoS One, 2015. 10(10): p. e0137703.

42. Ren, B., et al., E2F integrates cell cycle progression with DNA repair, replication, and G(2)/M checkpoints. Genes Dev, 2002. 16(2): p. 245–56.

43. Yu, H., Z. Li, and M. Wang, Expression and prognostic role of E2F transcription factors in high-grade glioma. CNS Neurosci Ther, 2020. 26(7): p. 741–753.

44. Patil, A.A., et al., FANCD2 re-expression is associated with glioma grade and chemical inhibition of the Fanconi Anaemia pathway sensitises gliomas to chemotherapeutic agents. Oncotarget, 2014. 5(15): p. 6414–24.

45. Lu, X., et al., Dysregulation of TFDP1 and of the cell cycle pathway in high-grade glioblastoma multiforme: a bioinformatic analysis. Genet Mol Res, 2016. 15(2).

46. Greco, C., et al., Development of Pyrazolo[3,4-d]pyrimidine Kinase Inhibitors as Potential Clinical Candidates for Glioblastoma Multiforme. ACS Med Chem Lett, 2020. 11(5): p. 657–663.

47. Velasco, G., Sanchez, C, Guzman, M, Towards the use of cannabinoids as antitumour agents. nature Reviews Cancer, 2012. 12: p. 436–444.

48. Fraguas-Sanchez, A.I., C. Martin-Sabroso, and A.I. Torres-Suarez, Insights into the effects of the endocannabinoid system in cancer: a review. Br J Pharmacol, 2018. 175(13): p. 2566–2580.

49. Petrecca, K., et al., Failure pattern following complete resection plus radiotherapy and temozolomide is at the resection margin in patients with glioblastoma. J Neurooncol, 2013. 111(1): p. 19–23.

50. Zhang, Y., et al., The p53 Pathway in Glioblastoma. Cancers (Basel), 2018. 10(9).

51. Cheng, L., et al., Identification of an IFN-beta-associated gene signature for the prediction of overall survival among glioblastoma patients. Ann Transl Med, 2021. 9(11): p. 925.

52. Ballou, Y., et al., 5-HT serotonin receptors modulate mitogenic signaling and impact tumor cell viability. Mol Clin Oncol, 2018. 9(3): p. 243–254.

53. Kachler, K., et al., Enhanced Acid Sphingomyelinase Activity Drives Immune Evasion and Tumor Growth in Non-Small Cell Lung Carcinoma. Cancer Res, 2017. 77(21): p. 5963–5976.

54. Bi, J., et al., Targeting glioblastoma signaling and metabolism with a re-purposed brain-penetrant drug. Cell Rep, 2021. 37(5): p. 109957.

55. Asada, M., et al., Depletion of serotonin and selective inhibition of 2B receptor suppressed tumor angiogenesis by inhibiting endothelial nitric oxide synthase and extracellular signal-regulated kinase 1/2 phosphorylation. Neoplasia, 2009. 11(4): p. 408–17.

56. Peters, M.A.M., et al., Serotonin and Dopamine Receptor Expression in Solid Tumours Including Rare Cancers. Pathol Oncol Res, 2020. 26(3): p. 1539–1547.

