## Supplementary material for "Transcription Profile And Pathway Analysis Of The Endocannabinoid Receptor Inverse Agonist AM630 In The Core And Infiltrative Boundary Of Human Glioblastoma Cells": SuppementaryTable S1 &S2

**Supplementary Data**

**Table S1.** The top regulated genes under AM630 treatment in the core and invasive margin cell populations. The core cell response is more significant with the up regulation of 1672 and down regulation of 2424 genes at the two-fold significant level, shown at left. The response of the invasive cell pouplation is more muted with only 815 up and 660 down regulated genes at the two fold cut-off.

**CORE cell AM630 response**

| GENE | Log(FC) | Z score | GENE | Log(FC) | Z score |
|---|---|---|---|---|---|
| DHRS2 | 7.5 | 7.55 | CXCR4 | -5.25 | -7.74 |
| FAM153A | 6.58 | 5.28 | CNTN1 | -4.93 | -5.78 |
| RSAD2 | 6.49 | 4.11 | CDH11 | -4.88 | -6.36 |
| GCM1 | 5.92 | 4.94 | PRRX1 | -4.67 | -6.54 |
| HTN3 | 5.8 | 9.14 | HAS2 | -4.66 | -5.49 |
| NR4A3 | 5.41 | 4.44 | NTM | -4.53 | -5.48 |
| ERV3-2 | 5.19 | 7.11 | SORBS2 | -4.53 | -7.39 |
| MYCT1 | 5.04 | 6.1 | WDR76 | -4.22 | -5.39 |
| SESN2 | 5.04 | 4.86 | LMO3 | -4.15 | -4.94 |
| LINC01419 | 4.97 | 5.18 | DAB1 | -4.11 | -6.69 |
| CMPK2 | 4.92 | 3.96 | TM4SF18 | -4.1 | -5.71 |
| TMEM106A | 4.9 | 4.15 | ANKS1B | -4.06 | -6.63 |
| PCDH17 | 4.82 | 4.03 | DKK1 | -4.05 | -4.81 |
| PDK4 | 4.79 | 4.29 | UBA6 | -4.05 | -6.54 |
| GLS | 4.72 | 4.98 | KMO | -4.04 | -4.71 |
| ARAP2 | 4.65 | 5.96 | POLE2 | -4.04 | -5.62 |
| IFIT1 | 4.64 | 3.8 | ARGLU1 | -3.95 | -5.51 |
| TLL2 | 4.6 | 6.13 | CENPU | -3.92 | -6.82 |
| ATF3 | 4.57 | 5.87 | MCMBP | -3.89 | -5.38 |
| ZCCHC2 | 4.48 | 5 | KCNMA1 | -3.87 | -5.65 |
| SERPINI1 | 4.47 | 4.98 | NRG1 | -3.86 | -5.18 |
| SEMA6A | 4.45 | 4.72 | WNT5A | -3.86 | -4.87 |
| FAM46C | 4.42 | 5.05 | RHOJ | -3.84 | -8.46 |
| CFAP43 | 4.32 | 5.32 | LDB2 | -3.83 | -4.74 |
| STK35 | 4.27 | 4.06 | CDC7 | -3.78 | -4.14 |

**Invasive margin cell AM630 response**

| GENE | Log(FC) | Z score | GENE | Log(FC) | Z score |
|---|---|---|---|---|---|
| INPP4A | 5.63 | 5.14 | MINK1 | -3.81 | -3.43 |
| SZRD1 | 5.05 | 3.86 | ATP6V0D2 | -3.63 | -2.77 |
| RAB1A | 4.78 | 5.73 | HDAC9 | -3.28 | -2.33 |
| PSMB4 | 4.36 | 3.45 | RSPH10B2 | -3.17 | -2.46 |
| DERL1 | 4.23 | 2.98 | SNHG4 | -2.94 | -2.23 |
| ASF1A | 3.92 | 2.99 | MACF1 | -2.81 | -3.01 |
| RPS16 | 3.75 | 3.06 | PRR11 | -2.7 | -2.82 |
| SMURF2 | 3.74 | 3.57 | QKI | -2.66 | -2.39 |
| H3F3AP4 | 3.66 | 2.9 | LY9 | -2.64 | -2.51 |
| PTP4A1 | 3.65 | 3.05 | NPAS2 | -2.59 | -5.12 |
| SAP30L | 3.51 | 4.85 | PPP3CB | -2.57 | -2.8 |
| CTDSPL2 | 3.48 | 4.16 | RGS12 | -2.49 | -2.57 |
| RPLP1 | 3.35 | 1.96 | HNRNPA3 | -2.47 | -3.47 |
| PAFAH1B1 | 3.34 | 2.28 | PCDHGA1 | -2.47 | -2.64 |
| SOD1 | 3.21 | 3.17 | ZEB2 | -2.46 | -5.2 |
| PIAS2 | 3.2 | -2.67 | THRA | -2.42 | -2.71 |
| MBNL2 | 3.17 | 4.55 | PPRC1 | -2.37 | -2.84 |
| UBE2M | 3.17 | 2.75 | SHCBP1L | -2.37 | -4.03 |
| SNORD68 | 3.14 | 2.14 | MFN2 | -2.36 | -2.95 |
| LAMTOR5 | 3.11 | 2.95 | SLC44A1 | -2.36 | -2.86 |
| MT2A | 3.07 | 3.95 | ATP6V0E1 | -2.35 | -3.43 |
| ETF1 | 3.04 | 1.94 | FGD2 | -2.3 | -3.4 |
| BUD31 | 3 | 3.44 | CDH2 | -2.26 | -2.93 |
| EIF5 | 2.99 | 3.68 | SETD8 | -2.26 | -3.27 |
| HIST1H2BD | 2.96 | 3.24 | HNRNPC | -2.22 | -2.3 |

**Table S2.** Drug-like compound expression profiles that correlate with AM630 driven expression changes in common between the core and invasive margin cells. The AM630 response in the invasive margin population is modest relative to that in the core population and does not return significant correlations with CMAP profiles. A composite profile constituting genes with combined Z scores five standard deviations away from the null consisting of 131 up and 111 down regulated genes has a similar CMAP analysis correlating drug profile with serotonin receptor antagonists also returning high scores.

| DRUG | SAME | OPPOSITE | prob |
|---|---|---|---|
| withaferin A | 96 | 16 | 4.34E-15 |
| parthenolide | 89 | 20 | 3.35E-12 |
| 15-delta prostaglandin J2 | 85 | 18 | 5.75E-12 |
| thioridazine | 103 | 30 | 1.04E-10 |
| fendiline | 57 | 10 | 4.77E-10 |
| fluphenazine | 66 | 15 | 9.98E-10 |
| LY-294002 | 119 | 51 | 2.45E-08 |
| lomustine | 70 | 19 | 3.02E-08 |
| mefloquine | 59 | 12 | 5.18E-08 |
| anisomycin | 96 | 33 | 8.92E-08 |
| astemizole | 69 | 20 | 1.63E-07 |
| calmidazolium | 61 | 17 | 1.76E-07 |
| vorinostat | 97 | 37 | 1.98E-07 |
| etacrynic acid | 45 | 9 | 6.01E-07 |
| semustine | 68 | 22 | 1.07E-06 |
| lanatoside C | 82 | 29 | 1.40E-06 |
| nortriptyline | 53 | 15 | 1.62E-06 |
| trichostatin A | 129 | 63 | 2.66E-06 |
| prochlorperazine | 71 | 25 | 2.90E-06 |
| terfenadine | 70 | 25 | 3.60E-06 |
| disulfiram | 43 | 8 | 4.34E-06 |
| proscillaridin | 74 | 31 | 5.36E-06 |
| trifluoperazine | 76 | 31 | 5.50E-06 |
| puromycin | 72 | 26 | 5.64E-06 |
| niclosamide | 50 | 14 | 6.18E-06 |
